# Shared genetic basis for brain structure, insulin resistance and inflammation in schizophrenia: a colocalization study

**DOI:** 10.1101/2025.07.23.666268

**Authors:** Amalie C. M. Couch, Benjamin I. Perry, Daniel B. Rosoff, Edward R. Palmer, Jack Rogers, David Ray, Rachel Upthegrove

## Abstract

Schizophrenia (SZ) is accompanied by structural brain alterations and elevated cardiometabolic risk, potentially linked through shared immune–metabolic mechanisms. To test for shared genetic underpinnings, we performed multi-trait colocalization across SZ, structural MRI measures (bilateral cortical thickness, grey matter volume), insulin resistance markers (TG:HDL-C ratio, fasting insulin adjusted for BMI), and IL-6 signalling traits (IL-6, IL-6R, IL-6ST) at 185 genome-wide significant loci. Colocalization was detected at 53 loci, including a robust cluster at the *SLC39A8* missense variant rs13107325 linking SZ with cortical structure, peripheral IL-6ST, and TG:HDL-C (PP_coloc_ = 0.73–0.99). A secondary, lower-confidence colocalization signal at *FGF21* (PP_coloc_ = 0.6) further implicated metabolic– behavioural coupling as a convergent pathway. Downstream analyses of rs13107325 pQTLs indicated enrichment in synaptic development and cardiometabolic pathways. Together, these findings identify rs13107325 as a shared causal variant bridging neurodevelopment, immunity, and metabolism, and highlight metal-ion transport and FGF21 signalling as potential therapeutic entry points to address the intertwined psychiatric and metabolic burden of SZ.

## Introduction

Individuals with schizophrenia (SZ) life expectancy 15 to 20 years shorter than the general population^1–4^, largely due elevated rates of metabolic and cardiovascular diseases^5^. While these physical health conditions have traditionally been viewed as consequences of medication or lifestyle, growing evidence suggests they may reflect shared biological mechanisms, particularly chronic low-grade inflammation^6^. SZ itself is a highly heritable, polygenic disorder shaped by numerous common variants of small effect and rarer variants of larger effect^7,8^, and it is possible that these have both central and peripheral effects, with impact on psychiatric and physical comorbidities reflecting their systemic nature.

Indeed, growing evidence suggests dysfunctional metabolism and inflammation are common pathological mechanisms across multiple organ systems, including the brain, in individuals with psychotic disorders^9^. Evidence includes insulin resistance (IR), which is detectable in up to 40% of individuals at the onset of psychosis, prior to medication exposure^10–12^ and increased low grade, chronic inflammation is prominently identified in unmedicated first-episode psychosis (FEP)^13–16^. Meta-analytic evidence clearly identifies that circulating concentrations of interleukin (IL)-6 are elevated in individuals with SZ^13,17^. IL-6 is a pleiotropic cytokine that signals through two distinct pathways: classical (cis) signalling, mediated by membrane-bound IL-6 receptor (IL-6R), and non-classical trans-signalling, when soluble (s)IL-6R engages the ubiquitously expressed co-receptor IL-6ST^18–20^. Genetically predicted levels of IL-6 have been previously linked with changes to 33 structural magnetic resonance imaging (sMRI) measures of GMV and cortical thickness (CT) ^21^, and robust evidence demonstrates changes to the same brain structures in individuals with SZ; decreased grey matter volume (GMV)^22^ is present at illness onset, while progressive cortical thinning is implicated through the course of the illness^23^.

Brain structure is both heritable and influenced by peripheral mechanism during development^24^. Human and animal studies indicate that prenatal immune activation, particularly elevated IL-6 levels, can disrupt neurodevelopment and increase vulnerability to psychosis in adulthood^25–29^. Previous colocalization studies have provided strong evidence for shared causal variants among type 2 diabetes mellitus (T2DM), body mass index (BMI), and schizophrenia (SZ), particularly within brain-derived neurotrophic factor (BDNF)–related pathways^30^. However, the extent to which peripheral and central traits, including brain structure, share a common genetic architecture remains unclear. This question is particularly relevant given the frequent co-occurrence of SZ with metabolic dysfunction, elevated peripheral IL-6 signalling, and alterations in grey matter volume (GMV) and cortical thickness (CT). We therefore hypothesize the existence of shared causal variants spanning both peripheral and central domains.

Therefore, the primary objective of this study was to identify shared causal variants between both peripheral and central phenotypes associated with schizophrenia. To investigate this, we applied hypothesis prioritisation for multi-trait colocalization (HyPrColoc)^31^, a Bayesian method for identifying shared causal variants across traits to identify specific loci where genetic variation may simultaneously impact systems. By extending previous colocalization analysis^30^ to include larger, more contemporary GWAS cohorts and brain structural traits, we were able to identify convergent risk loci across both peripheral and central domains. This application of novel multi-trait methodology provides a template for addressing the complex genetic pleiotropy underlying psychosis multimorbidity and for identifying shared biological mechanisms that may inform new therapeutic strategies.

## Methods

### Summary Statistics for Schizophrenia, Brain Structure, Insulin Resistance and IL-6 Signalling-related Traits

For SZ, we used publicly available summary data from the most recent Psychiatric Genomics Consortium (PGC) GWAS (76,755 cases and 243,649 controls)^8^. We also used publicly available summary GWAS data from large-scale consortia (Supplementary Table 1) for 8 well characterised traits: three structural MRI phenotypes (left hemisphere cortical thickness [LH-CT], right hemisphere cortical thickness [RH-CT], and normalised grey matter volume [GMV] for head size)^32^; two biomarkers of IR (TG:HDL-C and fasting insulin [FI] adjusted for BMI)^33,34^ and three IL-6 signalling proteins (IL-6, IL-6R, and IL-6ST)^35^. All GWAS were conducted in European adult samples and adjusted for population stratification, age, and sex, other than the brain imaging phenotypes which were adjusted for age, age^2^, sex, age×sex, and age^2^×sex. Ethical approval was obtained by the original GWAS authors in accordance with the protocols of each individual study.

### Multi-trait colocalization

We employed hypothesis prioritisation for multi-trait colocalization (HyPrColoc)^31^ to identify shared genomic risk loci (Supplementary Figure 1). HyPrColoc (R package *jrs95/hyprcoloc*, version 0.0.2) uses a Bayesian framework to estimate the posterior probability that multiple traits share a single causal variant by evaluating all possible causal configurations, thereby identifying a candidate variant without inferring the direction of association. Unlike conventional bivariate colocalization approaches, HyPrColoc allows the simultaneous assessment of multiple correlated traits, an essential advantage when investigating complex multimorbid relationships such as those linking psychiatric, metabolic, and inflammatory domains. This framework was chosen as it accommodates linkage disequilibrium structure, operates on GWAS summary statistics without requiring individual-level data, and enables the prioritisation of loci most likely to underlie shared biological pathways across traits^31^. Consequently, assumptions of data normality or homoscedasticity and tests of equal variance between groups are not required to be tested or explicitly accounted for in this analysis.

We first identified 185 unique lead index SNPs that demonstrated Bonferroni-significant evidence of locus-level correlation (GWAS *p* < 5×10^−8^) with brain structure, IR or IL-6 signalling-related traits (**Supplementary Table 1 and 2)**. To quantify colocalization of each trait, we generated “lookup” files in python (version 3.9.6) that included all SNPs located within ±500 kb of the lead index SNP. HyPrColoc was subsequently run in R (version 4.4.3) to detect shared genetic risk loci among trait clusters that included SZ. The major histocompatibility complex (MHC) region was excluded from all analyses owing to its high linkage disequilibrium, which impedes reliable identification of causal variants.

According to HyPrColoc^31^, the first prior (*p*) reflects the probability that at least one trait has a causal variant in the region, while the second prior (*p*_c_) represents the conditional probability that a variant is causal for a second trait, given it is causal for one trait^31^. The regional and alignment thresholds specify how closely association signals must overlap spatially and in pattern to be considered colocalised, thereby refining confidence in shared causal variants. Accordingly, our primary analysis employed the default variant-specific priors (*p* = 1×10^−4^; *p*_c_ = 0.05) along with regional and alignment thresholds set at 0.5 to identify all colocalizing trait clusters across the 185 lead SNP loci. UpSet (R package *UpSetR* version 1.4.0) plots were then generated to visualize these clusters.

### Sensitivity Analysis

Using HyPrColoc’s Bayesian framework allows us to formally incorporate prior probabilities and quantify the strength of evidence for shared genetic signals across traits by varying *p*_c_, regional and alignment thresholds, thereby enhancing the interpretability. Therefore, to evaluate the robustness of colocalization signals and cluster stability, we repeated the analysis using both increasingly stringent *p*_c_ values (0.05, 0.02, 0.01, 0.005, 0.001) and increasingly stringent regional and alignment thresholds (0.5, 0.6, 0.7, 0.8, 0.9). Cluster frequency plots and line graphs of posterior probabilities across increasing *p*_c_ thresholds were generated for each cluster containing schizophrenia (SZ) among the colocalizing traits. Where evidence of potential colocalization with SZ was observed, heatmaps based on a cluster similarity matrix were created to visualize cluster stability across threshold permutations. Additionally, stacked regional association plots were produced to inspect candidate SNPs, forest plots to assess the strength of their associations across traits, and the local linkage disequilibrium (LD) structure within the genomic regions.

### Functional Characterisation in Silico of Candidate SNPs

#### Quantitative Trait Loci Analysis

To investigate the functional impact of candidate SNPs that colocalized SZ with brain structure, insulin resistance and inflammation traits, we investigated whether candidate SNPs were quantitative trait loci (QTL), focusing on expression QTLs (eQTLs) and protein QTLs (pQTLs). eQTL data were sourced from the GTEx Portal (version 8), using the package *gtexr* (version 0.2.1)^36^, to identify tissue-specific gene expression data associated with candidate SNPs across all human tissues for which data was available in GTEx. pQTL data were obtained from the UK Biobank Pharma Proteomics Project (UKB-PPP), which uses aptamer-based proteomic profiling of circulating protein levels in approximately 50,000 individuals from the UK Biobank^35^. Up- and down-regulated pQTLs (association *p* < 1.7×10^−11^) as a result of candidate variants were extracted for pathway analyses.

#### Protein-Phenotype Associations

To complement the pQTL findings, we additionally examined the epidemiological associations of the corresponding proteins with incident schizophrenia diagnoses in the UK Biobank. Protein–disease associations were obtained from the *Atlas of the Plasma Proteome in Health and Disease* (https://proteome-phenome-atlas.com/); a large-scale proteome–phenome resource integrating plasma proteomics with disease outcomes in 53,026 participants ^*37*^. We extracted effect estimates for the endpoint *“Schizophrenia or delusion, excluding more controls”* based on Cox proportional hazards models reported in the atlas, providing longitudinal evidence of protein–disease associations. This lookup enabled evaluation of whether genetically implicated proteins from our pQTL analyses also show observational associations with schizophrenia incidence in the general population.

#### Pathway Enrichment Analysis

To characterise biological pathways linked to SNP-associated pQTLs, we then conducted enrichment analyses using the *EnrichR* package (version 3.4). Over-representation analysis (ORA) was performed separately for proteins exhibiting increased or decreased expression correlated with the minor allele. Gene Ontology (GO) Biological Process and Kyoto Encyclopaedia of Genes and Genomes (KEGG) pathway databases were used for annotation. Enrichment significance was assessed using gene ratio metrics and adjusted p-values using the BH method, then sub-setting the top 10 significant pathways (FDR < 5%, ranked by Fold Enrichment). Additionally, gene set enrichment analysis (GSEA) was carried out on a ranked list of all pQTL-associated genes. GSEA employed three curated databases: DisGeNET (gene-disease associations) and MSigDB Hallmark (50 canonical gene sets). Statistical significance in GSEA was evaluated using odds ratios and adjusted p-values, with significant gene sets (BH correction FDR < 5%) plotted.

#### Expression Profiling in RNA Sequencing Datasets

To examine the developmental expression pattern of genes implicated at colocalised loci, we also analysed three publicly available transcriptomic datasets of the developing human brain. First, single-cell RNA sequencing data from Wang et al. (2025) were interrogated to assess gene expression across annotated cell populations in the developing human cortex at multiple stages during development, including vascular cells, glia, and neuronal subtypes^38^. Second, using the Gene Expression in Cortical Organoids (GECO) online tool, time-course bulk RNA-seq data from Gordon et al. (2021) of human cortical organoids were used to evaluate gene expression during in vitro differentiation^39,40^. Third, likewise using the GECO tool, bulk RNA-seq data from the BrainSpan Atlas of the Developing Human Brain were extracted^40^, spanning prenatal to adulthood stages (8 post-conception weeks to 40 years).

## Results

### Multi-trait colocalization between schizophrenia, cortical structure, biomarkers of insulin resistance and IL-6 signalling

#### Primary Analysis

Using default priors (PP_coloc_ ≥ 0.5; *p* = 1 x 10^−4^; *p*_*c*_ = 0.05; regional/alignment thresholds = 0.5), colocalizing trait clusters were identified at 53 of 185 lead SNP loci (Figure 1**A** and **Supplementary Table 3**). Five clusters involved SZ, (Figure 11B), four of which showed colocalization exclusively with TG:HDL-C. These included candidate variants in *RBM6* (rs6765484; intron; PP_coloc_ = 0.75), *STARD3* (rs881844; intron; PP_coloc_ = 0.68), *FGF21* (rs838133; synonymous; PP_coloc_ = 0.6041), and an intergenic locus between *RPL30P11 and GNAI2P1* (rs61920311; PP_coloc_ = 0.92) (Table 1). The fifth cluster demonstrated shared causality between SZ, IL-6ST, TG:HDL-C, bilateral cortical thickness, and grey matter volume at *SLC39A8* (rs13107325; missense; PP_coloc_ = 0.55) (Table 1). Of the five colocalizing candidate variants, two candidates serve as genome-wide significant SZ index SNPs (rs13107325 and rs61920311). Lead and candidate SNPs were identical in two cases (rs838133 and rs13107325) and distinct in the remaining three (Table 1).

**Figure 1:**
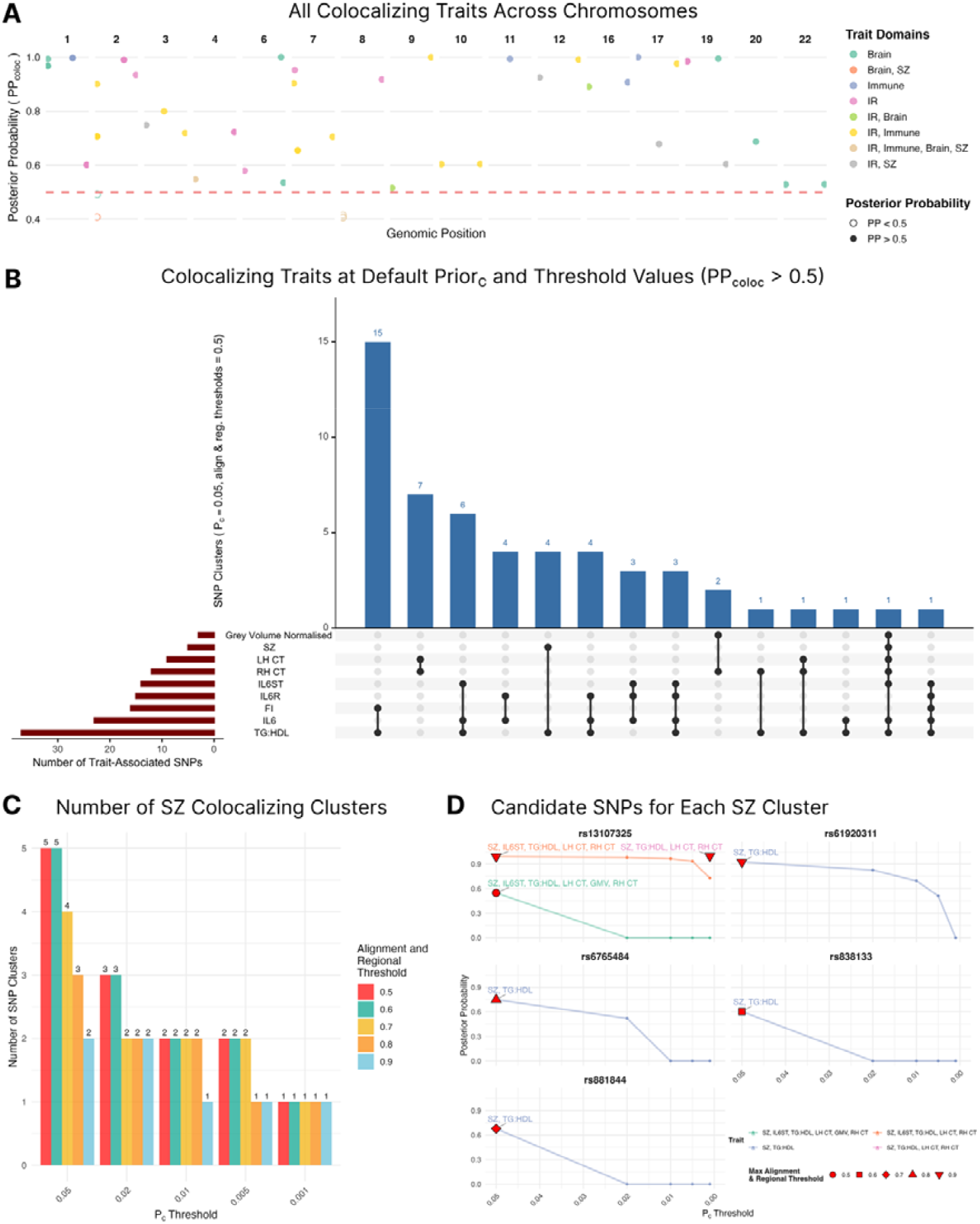
Colocalizing trait clusters identified at default priors and threshold values. (A) the genomic distribution of all colocalizing traits identified through HyPrColoc analysis at default settings across chromosomes. Each point represents a shared lead variant associated with a cluster of traits sharing a causal variant, colour-coded by trait domain combinations (Brain, Schizophrenia, Immune, or Insulin Resistance [IR]). The y-axis indicates the posterior probability of colocalization (PP_coloc_), with the dashed red line marking the significance threshold of PP_coloc_ > 0.5. Variants above this threshold indicate modest evidence for shared causal effects between schizophrenia and other trait domains. (B) UpSet plot summarising all colocalizing trait clusters with PP_coloc_ > 0.5 at default variant-specific priors (*p* = 1 × 10^−4^, *p*_*c*_ = 0.05) and regional/alignment thresholds (0.5). The blue bars indicate the number of SNP clusters for each trait combination, and the connected black dots indicate the traits included in each cluster. The horizontal red bars on the left show the total number of trait-associated SNPs per trait. (C) Number of SZ-colocalizing clusters across decreasing *p*_*c*_ thresholds (x-axis) and multiple alignment and regional thresholds (color-coded). Only clusters containing SZ are shown. (D) Posterior probabilities (PP_coloc_) of candidate shared casual SNPs for each SZ-containing cluster across *p*_*c*_ thresholds. Only SZ-associated clusters are shown. Points and lines indicate how PP_coloc_ changes with thresholding, and symbols indicate the maximum alignment and regional threshold at which each cluster was initially identified. Variation and uncertainty are quantified through PP_coloc_ derived from the Bayesian model, rather than conventional within-group variance.

**Table 1:** Primary results from HyPrColoc analysis, tabulating SNPs clustering traits with SZ at default settings (*p* = 1×10^−4^, *p*_c_ = 0.05, alignment and regional thresholds = 0.5). Mean Minor allele frequency (MAF) of the candidate SNP was derived from European reference population via LDLink in the 1000 Genomes dataset. PP_coloc_ represents the posterior probability that there is a single shared causal SNP amongst traits. PP_explained_ represents the proportion of posterior probability that is explained by the candidate SNP given. N SNPs are the number of SNPs present in all trait datasets compared. †SNP is an index SNP for schizophrenia with an association p < 5×10^−8^. * PP_coloc_ increased to >0.90 in sensitivity analysis after sensitivity analysis removed GMV from the cluster with increasing regional and alignment thresholds (see Sensitivity Analysis section).

| Lead SNP | Candidate SNP | Region<sub>candidate</sub> | MAF<sub>candidate</sub> | Gene Implicated | Variant Type | Colocalised Traits with SZ | $PP_{\text{coloc}}$ | $PP_{\text{explained}}$ | N SNPs |
| --- | --- | --- | --- | --- | --- | --- | --- | --- | --- |
| rs11130227 | rs6765484 | 3p21.31 | 0.46 | *RBM6* | Intron | TG:HDL-C | 0.75 | 0.04 | 1092 |
| rs931992 | rs881844 | 17q12 | 0.33 | *STARD3* | Intron | TG:HDL-C | 0.68 | 0.28 | 660 |
| rs838133 | rs838133 | 19q13.33 | 0.43 | *FGF21* | Synonymous | TG:HDL-C | 0.6041 | 1.00 | 2315 |
| rs17402950 | rs61920311† | 12p13.1 | 0.45 | *RPL30P11*,<br>*GNAI2P1* | Intergenic | TG:HDL-C | 0.92 | 0.37 | 2324 |
| rs13107325 | rs13107325† | 4q24 | 0.08 | *SLC39A8* | Missense | IL-6ST, TG:HDL-C,<br>LH-CT, RH-CT,<br>GMV | 0.55* | 1.00 | 2049 |

#### Sensitivity Analysis

As *p*_c_ decreased, SZ-containing clusters were progressively reduced from five (at *p*_c_ = 0.05) to one (*p*_c_ = 0.001) (Figure 1B-C). PP_coloc_ values for SZ-TG:HDL-C clusters at rs838133 and rs881844 declined sharply beyond *p*_c_ = 0.05 (Figure 1D). Similarly, the SZ-containing trait cluster at rs13107325 persisted only beyond *p*_c_ = 0.05 without GMV (Figure 1D). SZ-TG:HDL-C clusters at rs6765484 and rs61920311 were absent beyond *p*_c_ = 0.02 and *p*_c_ = 0.005, respectively (Figure 1D). Although the cluster of SZ, TG:HDL-C, bilateral cortical thickness and IL6ST shared the causal variant rs13107325 at *p*_c_ = 0.001 (PP_coloc_ = 0.73; alignment and regional thresholds = 0.8), increasing alignment/regional thresholds to 0.9 yielded a higher posterior probability for SZ, TG:HDL-C, and cortical thickness (PP_coloc_ = 1.0), suggesting a tightly aligned shared genetic architecture (Figure 1D).

### Characterisation of rs13107325

Given its stability across sensitivity thresholds and inclusion of brain, metabolic, and immune domains (**Supplementary Figure 2**), rs13107325 was prioritised for functional characterisation. **eQTL and pQTL Analyses** rs13107325 did not meet FDR significance for *SLC39A8* expression in GTEx (FDR>0.05) (Figure 2**A**). At nominal thresholds, the minor allele was associated with reduced *SLC39A8* expression in blood, liver, esophagus muscularis, spinal cord, and nucleus accumbens, and increased expression in breast, salivary gland, and ileum tissues (*p* < 0.05; FDR ≈ 0.23). It was a significant eQTL only for PABPC1P7 in cerebellum (*p* = 1.93×10^−4^; NES = 0.62). Although PABPC1P7 is a pseudogene, the PABPC1 family regulates mRNA translation and stability, suggesting potential regulatory relevance.

**Figure 2:**
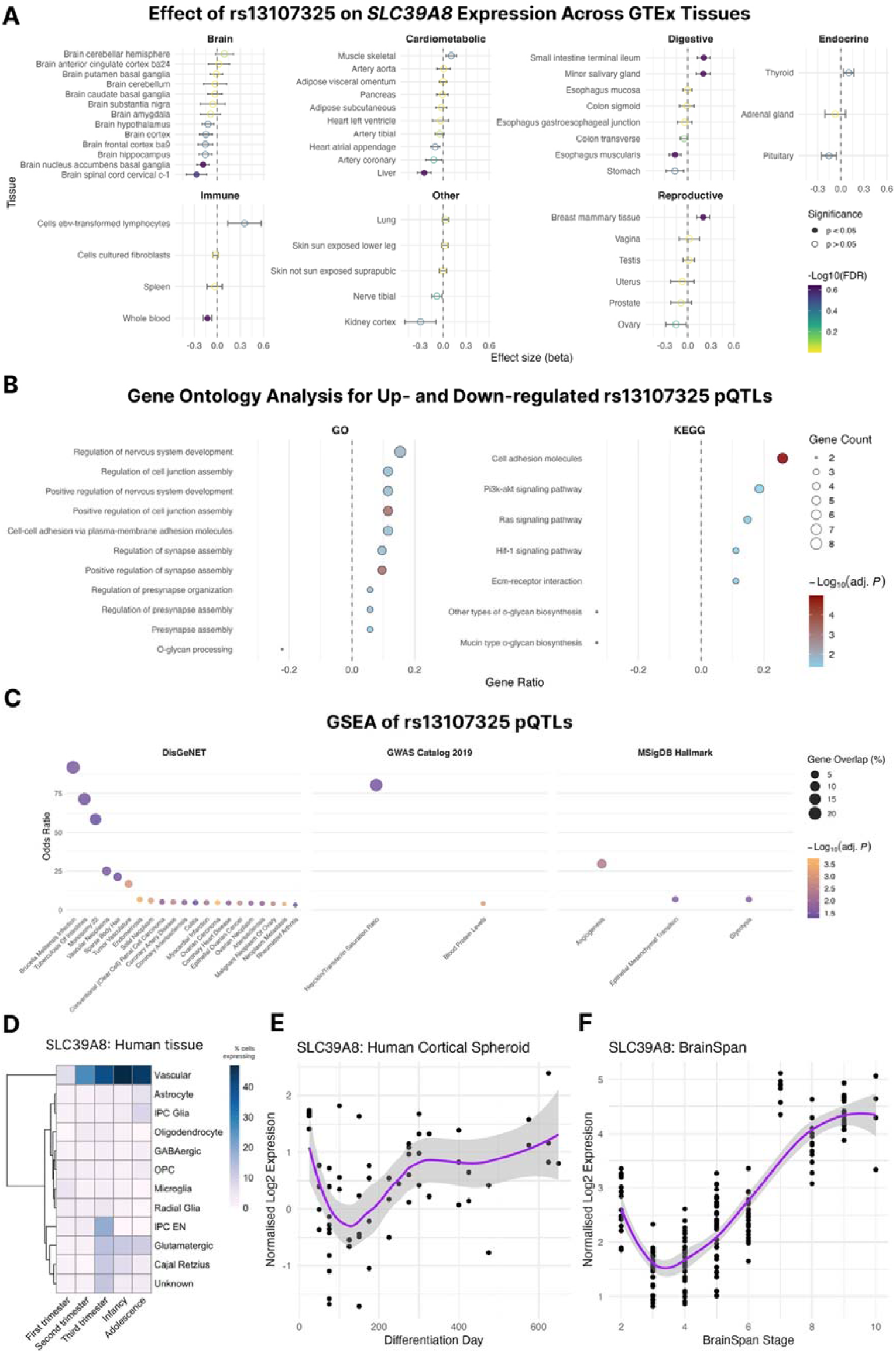
Pathway analysis of rs13107325 pQTLs and developmental expression of ZIP8. (A) Effect (B) Over-representation analysis (ORA) of genes whose protein levels are associated with the rs13107325 variant. GO biological process terms (left panels) and KEGG pathways (right panels) are shown for proteins upregulated (top) or downregulated (bottom) with the minor allele. Dot size represents gene ratio; colour represents adjusted p-value. (C) Gene set enrichment analysis (GSEA) of rs13107325 pQTLs using DisGeNET, GWAS Catalog, and MSigDB hallmark gene sets. Dot size reflects gene overlap percentage; colour reflects adjusted p-value; x-axis indicates odds ratio. (D) Heatmap showing percentage of cells expressing ZIP8 across different human brain cell types and developmental stages, data from publicly available scRNA-seq ^38^. ZIP8 is most strongly expressed in vascular cells, peaking in infancy. (E) Temporal expression profile of ZIP8 in human cortical spheroids across differentiation time, with data from publicly available bulk-RNAseq ^39^. Data represent normalized log2 expression values with a LOESS regression fit and 95% confidence interval. (F) ZIP8 expression in the human brain across developmental stages in the BrainSpan dataset ^40^; RNA sequencing and exon microarray data profiling of up to sixteen cortical and subcortical structures across the full course of human brain development. Developmental stages are: early prenatal (8–10 post-conceptional weeks [PCW], Stage 2), mid prenatal (10–13 PCW, Stage 3; 13–16 PCW, Stage 4; 16–19 PCW, Stage 5; 19–24 PCW, Stage 6), late prenatal (24–38 PCW, Stage 7), early infancy (birth–6 months, Stage 8), late infancy (6–19 months, Stage 9), early childhood (19 months–5 years, Stage 10), middle childhood (6–11 years, Stage 11), adolescence (12–19 years, Stage 12), and young adulthood (21–40 years, Stage 13). Expression values are normalized log_2_ values with LOESS smoothing and 95% confidence interval.

pQTL analysis identified 64 proteins significantly associated with rs13107325 (*p* < 1.7×10^−11^) (**Supplementary Figure 3, Supplementary Table 4**). Upregulated proteins were enriched for synapse assembly and regulation of nervous system development (GO enrichment: pre-synapse, FE = 30.4; synapse assembly, FE = 10.9; p-adj = 0.018). KEGG pathways implicated cell adhesion molecules, ECM–receptor interactions, and PI3K–AKT, HIF-1, and Ras signalling. Downregulated proteins were enriched for O-glycan processing (FE = 105.5; p-adj = 0.045) and biosynthesis pathways (p-adj = 3.7×10^−3^) (**Figure 2B**).

Gene set enrichment analysis (GSEA) of the 63 pQTLs identified enrichment for angiogenesis (OR = 29.7; p-adj = 0.004), glycolysis (OR = 6.7; p-adj = 0.024), and cardiometabolic traits such as coronary artery disease and myocardial infarction (p-adj < 0.01) (Figure 2C).

#### Protein–Phenotype Associations

In the UK Biobank Proteome–Phenome Atlas (∼43,000 participants; 183 incident SZ cases), 22 rs13107325-linked proteins were associated with incident SZ (*p* < 0.05; **Supplementary Figure 4**). Elevated circulating Folate receptor α (HR = 2.30), Agrin (HR = 1.98), and Angiopoietin-2 (HR = 1.86) predicted higher risk, while reduced TrkB (HR = 0.55), NELL1 (HR = 0.55), and GPRG2 (HR = 0.49) were associated with reduced risk. These findings link genetically anchored proteins to schizophrenia incidence, reinforcing angiogenic, glycolytic, and cardiometabolic mechanisms as shared pathways in SZ pathophysiology.

#### Developmental Expression

To determine the spatiotemporal context in which rs13107325 may exert its effects during development, we examined *SLC39A8* expression in publicly available single-cell RNA sequencing datasets from human foetal brain tissue, alongside in vitro differentiation models using human induced pluripotent stem cell (hiPSC)-derived cortical organoids and the BrainSpan dataset.

Single-cell and bulk transcriptomic analyses showed highest *SLC39A8* expression in vascular cell types of the developing brain, peaking in infancy (Figure 2D). In hiPSC-derived cortical organoids and BrainSpan datasets, expression increased steadily from embryogenesis to postnatal stages (Figure 2E-F). These findings suggest that rs13107325 may exert effects during critical neurodevelopmental windows, potentially linking neurovascular and synaptic maturation to later psychiatric and metabolic vulnerability.

## Discussion

Our multi-trait colocalization analysis identified rs13107325 in *SLC39A8* as a shared causal variant linking schizophrenia, TG:HDL-C, bilateral cortical thickness, and plasma IL-6ST.

Additional loci, including *RBM6, STARD3*, and *FGF21*, showed shared effects between SZ and lipid traits, suggesting broader cross-system coupling between brain structure, metabolism, and inflammation.

This study provides the first genomic evidence of a single variant jointly influencing neuroanatomical, metabolic, and cytokine domains within schizophrenia, extending previous reports implicating *SLC39A8* in HDL, TG, BMI, and type 2 diabetes^30^. The missense variant rs13107325 (C/T; p.Ala391Thr) disrupts the divalent metal ion transporter ZIP8, affecting manganese and zinc transport^41^. Functional studies demonstrate that this loss-of-function allele alters metal homeostasis, reduces dendritic spine density, impairs synaptic glutamatergic transmission, and compromises blood–brain barrier integrity^42–45^. These mechanistic effects align with human neuroimaging findings linking the risk allele to altered basal ganglia tissue composition^42^ and cognitive performance^43^. Collectively, these data position rs13107325 as a pleiotropic hub at the intersection of neurodevelopment, immunity, and metabolism.

Our functional and proteomic analyses reinforce this model. The rs13107325-linked proteome was enriched for pathways governing synapse assembly, angiogenesis, and glycolysis, consistent with prior evidence that ZIP8 regulates both neuronal and endothelial homeostasis^45,46^. Developmental expression data revealed that *SLC39A8* is predominantly expressed in brain vascular cells, increasing from foetal to postnatal life, highlighting a developmental period during which metal transport disruptions could influence neurovascular maturation. Moreover, rs13107325 pQTLs predicted incident SZ in population-level data, with notable effects of Folate receptor α—previously linked to disrupted one-carbon metabolism and neurotransmitter synthesis^47,48^. Together, these convergent results implicate micronutrient transport, vascular metabolism, and synaptic development as interconnected biological axes shaping schizophrenia risk.

Although colocalization between SZ and TG:HDL-C at *FGF21* was less stable across priors, its biological relevance warrants emphasis. *FGF21* encodes a hepatokine central to metabolic adaptation, with distinct variants regulating circulating FGF21 and macronutrient preference ^49^. Experimental and clinical studies show FGF21 agonism improves insulin sensitivity, reduces inflammation, and decreases alcohol consumption^49–51^. Our findings suggest that *FGF21* may represent a genetic nexus linking insulin resistance, diet-related behaviour, and neural pathways relevant to SZ, offering a tractable therapeutic target. Given that several FGF21 analogues (e.g., pegozafermin, efruxifermin) are in late-stage trials for metabolic disease^52^, evaluating their effects in psychiatric populations could illuminate shared therapeutic mechanisms across metabolic and cognitive domains.

While leveraging large, well-powered GWAS datasets and high-throughput colocalization enhances confidence in shared loci, several limitations remain. Colocalization cannot infer causal directionality; functional and Mendelian randomization studies are needed to determine whether immune–metabolic dysfunction drives brain alterations or vice versa. Proxy measures such as TG:HDL-C may incompletely capture insulin resistance biology, and pleiotropic signals may partly reflect linked loci. Additionally, sex-specific or ancestry-dependent effects were not assessed, limiting generalisability. Expanding analyses to include non-European cohorts and region-specific brain metrics will help clarify the scope of shared genetic architecture.

## Conclusion

Through integrative colocalization across neuroanatomical, metabolic, and inflammatory traits, we identify rs13107325 in *SLC39A8* as a shared causal variant connecting schizophrenia with immune–metabolic dysregulation and cortical structure. This missense variant disrupts metal-ion transport, offering a biologically coherent mechanism linking neurodevelopment, energy metabolism, and systemic inflammation. Complementary evidence at *FGF21* further highlights metabolic–behavioural coupling as a convergent pathophysiological axis. The identification of rs13107325 and its associated proteomic network offers actionable targets for intervention. Modulating ZIP8-mediated metal transport or pharmacologically activating FGF21 pathways could mitigate both psychiatric and metabolic dysfunction in high-risk individuals. Moreover, population-level protein biomarkers linked to this variant offer opportunities for early risk stratification and precision medicine approaches in schizophrenia, bridging molecular genetics with clinical decision-making. Together, these findings support therapeutic exploration of metal transport and FGF21 signalling pathways as potential means to address the intertwined psychiatric and cardiometabolic burden of schizophrenia.

## Acknowledgements

ACMC and RU are supported by the NIHR Oxford Health Biomedical Research Centre (NIHR203316). BIP is supported by an NIHR Advanced Fellowship (NIHR304365). The views expressed are those of the author(s) and not necessarily those of the NIHR or the Department of Health and Social Care. EP is supported by the Wellcome Trust Midlands Mental Health & Neurosciences PhD Programme for Healthcare Professionals. JR is supported by the University of Birmingham.

## Data and Code Availability

Summary statistics for schizophrenia (SZ) were obtained from Trubetskoy et al. (2022) (https://pgc.unc.edu/for-researchers/download-results/), cortical thickness and peripheral grey-matter volume from Smith et al. (2021) (https://open.oxcin.ox.ac.uk/ukbiobank/big40/), circulating IL-6, IL-6R, and IL-6ST levels from Sun et al. (2023) (https://www.synapse.org/Synapse:syn51364943/wiki/622119), fasted insulin adjusted for BMI from Lotta et al. (2016) (https://www.magicinvestigators.org/downloads/index.html), and triglyceride to HDL-C ratio (TG:HDL-C) from Oliveri et al. (2024) (https://www.ebi.ac.uk/gwas/studies/GCST90295949). All analysis scripts, including those implementing HyPrColoc (Foley et al., 2021) for multi-trait colocalization and downstream variant characterisation analysis, are available on GitHub at https://github.com/acmc96/hyprcoloc_SZ_IL6_sMRI_IR. For developmental gene expression analyses, publicly available transcriptomic datasets were used: single-cell RNA-seq from Wang et al. (2025) across annotated cortical cell populations (https://cell.ucsf.edu/snMultiome/), time-course bulk RNA-seq of human cortical organoids from Gordon et al. (2021) accessed via the GECO tool (http://solo.bmap.ucla.edu/shiny/GECO), and bulk RNA-seq from the BrainSpan Atlas of the Developing Human Brain (8 post-conception weeks to 40 years), also accessed via GECO (http://solo.bmap.ucla.edu/shiny/GECO).

