## Supplementary Figure for "Shared genetic basis for brain structure, insulin resistance and inflammation in schizophrenia: a colocalization study"

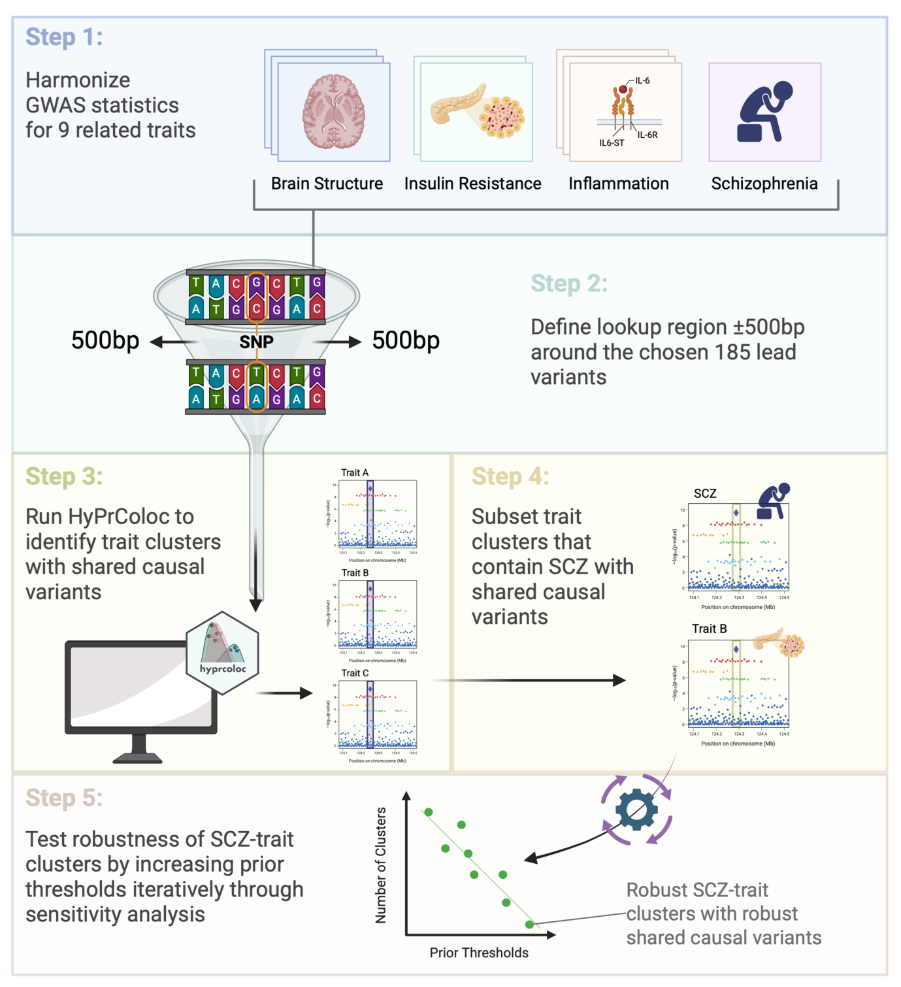

**Supplementary Figure 1: Workflow of HyPrColoc analysis to identify shared causal variants between schizophrenia and related traits.** This flowchart illustrates the analytical pipeline used to perform HyPrColoc (Hypothesis Prioritisation in multi-trait Colocalization) analysis ^31^ . Step 1: GWAS summary statistics for nine biologically related traits (SZ, peripheral IL-6R, IL-6ST, IL-6, GMV, Left/Right Hemisphere Cortical Thickness, Fasting Insulin and TG:HDL-C) across the four domains of interest (e.g., brain structure, insulin resistance, inflammation, SZ) were harmonized to ensure compatibility across datasets. Step 2: A lookup region of ±500 base pairs was defined at 185 lead variants (see **Supplementary Table 5**) of interest. Step 3: HyPrColoc was run to identify clusters of traits sharing causal variants. Step 4: Trait clusters containing schizophrenia and shared causal variants were subset for further examination. Step 5: The robustness of identified schizophrenia–trait clusters was tested through iterative sensitivity analyses by increasing prior_c_, alignment and regional thresholds to confirm stability of shared causal variant signals. Figure created with BioRender.

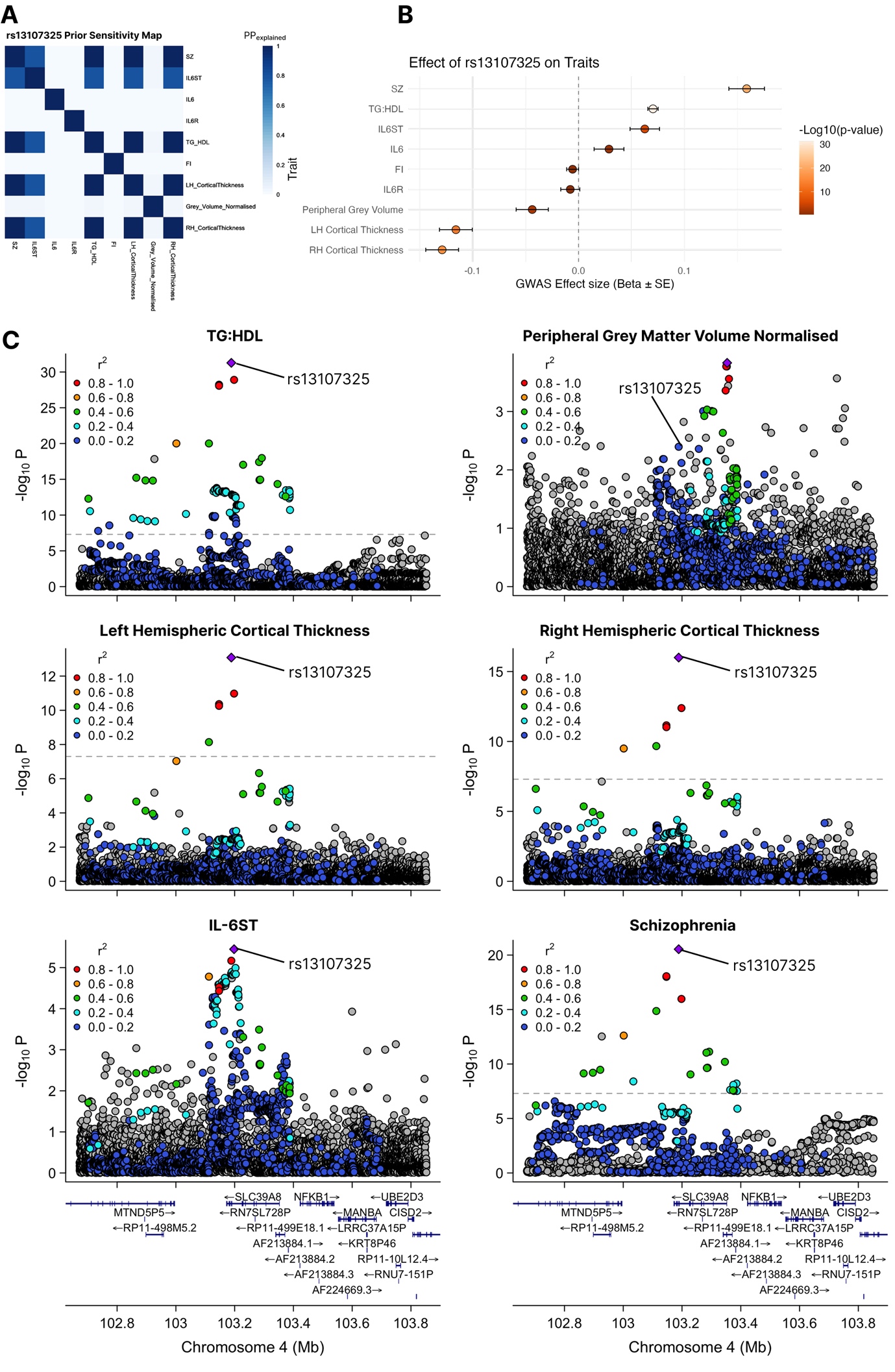

**Supplementary Figure 2: rs13107325 colocalization results.** (A) Sensitivity plot of candidate SNP rs13107325 across all traits, demonstrating repeated calls of the hyprcoloc function to compute a similarity matrix which illustrates how strongly clustered/colocalized pairs of traits are across different input thresholds and priors. The darker the blue, the more robust colocalization of trait pairs is found to be in sensitivity analyses. (B) Variant Effect Size forest plot in all traits used in the study, data from GWAS studies laid out in Table 1. (C) Regional genetic association plots for the most robust colocalised loci returning evidence for colocalization between schizophrenia, brain structure, insulin resistance and IL-6 signalling. Regional association plots denote chromosomal location (x axis) and strength of association with listed trait (−log10(p)) (y axis) alongside genomic position. SNP r^2^ estimated from the European panel from 1000 Genomes cohort. Variant rs13107325 labelled on each plot.

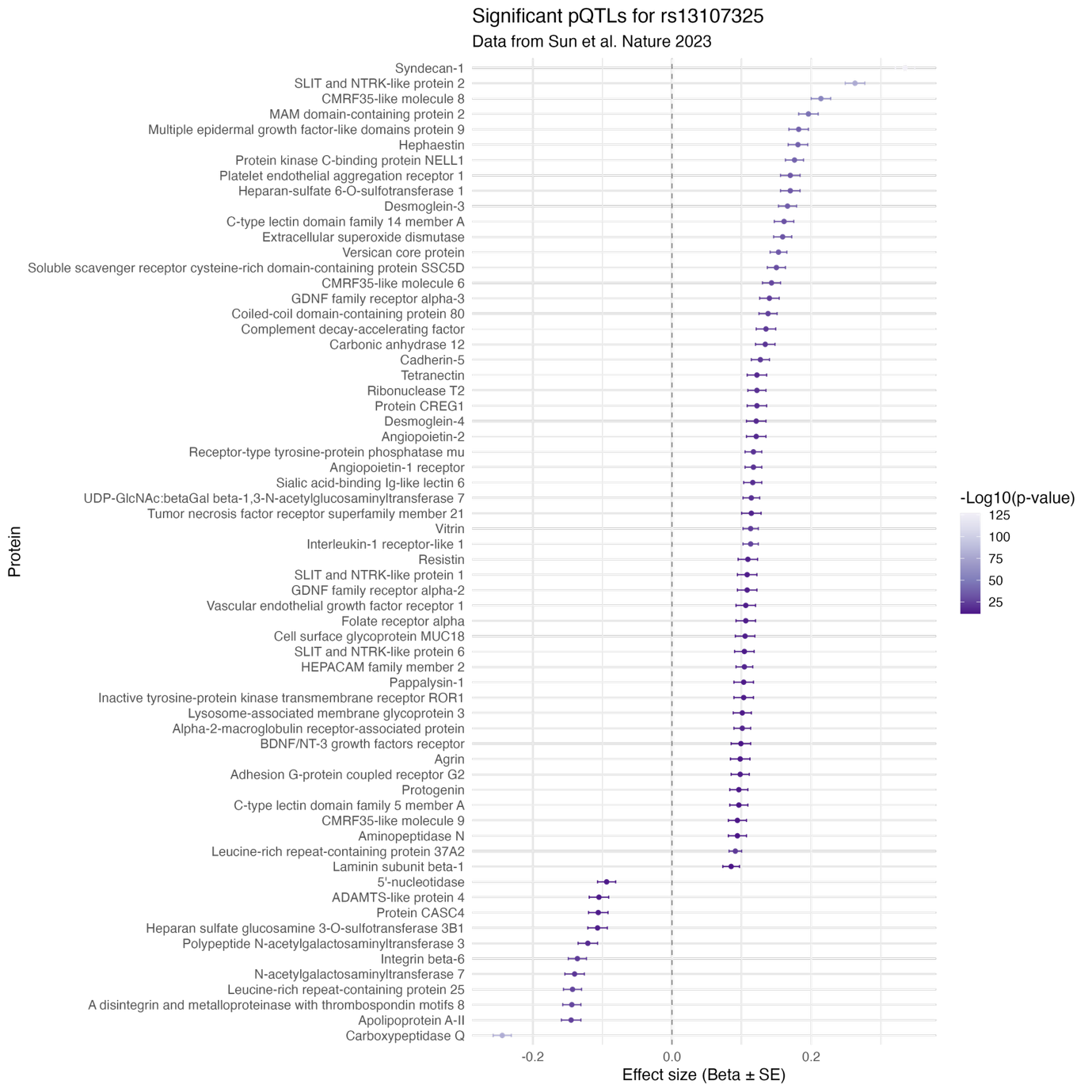

**Supplementary Figure 3: Forest plot of protein quantitative trait loci (pQTL) associations with rs13107325 (P < 1.7x10^-11^), with data derived from the UKB-PPP (29).** This forest plot displays standardized effect sizes (β coefficients) and 95% confidence intervals for the association between the schizophrenia risk SNP rs13107325 and plasma levels of individual proteins, based on pQTL data. Proteins are ranked on the y-axis by effect size, with positive associations shown to the right and negative associations to the left of the zero line. Effect size magnitudes and statistical significance are visualized using purple circles, with circle shading indicating –log_10_(P). The plot highlights proteins with diverse biological functions, including extracellular matrix components (e.g., Syndecan-1, Heparan sulphate 6-O-sulfotransferase), immune markers (e.g., Complement decay-accelerating factor, Interleukin-1 receptor-like 1), metabolic enzymes, and vascular/endothelial factors (e.g., Angiopoietin-like protein 2, VEGF receptor 1).

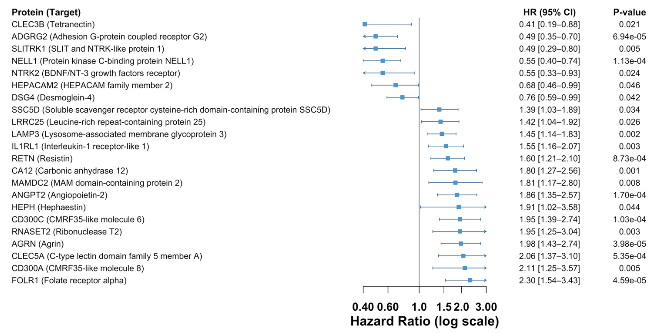

**Supplementary Figure 4:** Proteins influenced by the SLC39A8 missense variant (rs13107325) and risk of incident schizophrenia in UK Biobank. Forest plot of Cox proportional-hazards estimates for the top nominally significant (P < 0.05) circulating proteins whose levels are associated with rs13107325 and with subsequent schizophrenia diagnosis in UK Biobank. Points show hazard ratios (HR) per SD increase in protein level; horizontal bars indicate 95% CIs; the x-axis is on a log scale with the vertical line at HR = 1 (no association). Protein names are listed at left (gene symbol; common name), and nominal P-values are shown at right. HR < 1 indicates lower risk (putative protective association) and HR > 1 indicates higher risk.
